# The evolution of virulence in *Pseudomonas aeruginosa* during chronic wound infection

**DOI:** 10.1101/2020.05.29.124545

**Authors:** Jelly Vanderwoude, Derek Fleming, Sheyda Azimi, Urvish Trivedi, Kendra P. Rumbaugh, Stephen P. Diggle

## Abstract

Opportunistic pathogens are associated with a number of chronic human infections, yet the evolution of virulence in these organisms during chronic infection remains poorly understood. Here, we tested the evolution of virulence in the human opportunistic pathogen *Pseudomonas aeruginosa* in a murine chronic wound model using a two-part serial passage and sepsis experiment, and found that virulence evolved in different directions in each line of evolution. We also assessed *P. aeruginosa* adaptation to a chronic wound after 42 days of evolution, and found that morphological diversity in our evolved populations was limited compared to that previously described in cystic fibrosis (CF) infections. Using whole genome sequencing, we found that genes previously implicated in *P. aeruginosa* pathogenesis (*lasR, pilR, fleQ, rpoN*, and *pvcA*), acquired mutations during the course of evolution in wounds, with some mutations evolving in parallel across all lines of evolution. Our findings highlight that (i) *P. aeruginosa* heterogeneity may be less extensive in chronic wounds than in CF lungs; (ii) genes involved in *P. aeruginosa* pathogenesis acquire mutations during chronic wound infection; (iii) similar genetic adaptations are employed by *P. aeruginosa* across multiple infection environments, and (iv) current models of virulence may not adequately explain the diverging evolutionary trajectories observed in an opportunistic pathogen during chronic wound infection.

## INTRODUCTION

Opportunistic pathogens, those that only cause disease in hosts with compromised immune defenses, are responsible for several chronic, treatment-resistant infections in humans, such as certain skin, respiratory, and urinary tract infections. Common problematic human opportunists include *Pseudomonas aeruginosa, Staphylococcus aureus, Streptococcus pneumoniae, Candida albicans, Klebsiella pneumoniae, Serratia marcescens*, and *Acinetobacter baumannii*. While chronic infections caused by opportunists are prevalent in both community and hospital environments, the complex nature of their virulence remains elusive. Investigating the dynamics of virulence in chronic infections is of rising interest as researchers turn to novel treatments, such as anti-virulence drugs, to combat rapidly increasing antimicrobial resistance [1-4]. Yet, there remain two core questions for which the answers are unclear: (i) how does virulence typically evolve in opportunistic pathogens, and (ii) are these patterns of evolution predictable?

Currently, there are four classic hypotheses which explain how pathogenic virulence evolves, where virulence is attributed to either (i) new host-parasite associations; (ii) short-sighted evolution; (iii) evolutionary trade-offs; or (iv) coincidental selection [5-7]. The first of these, the ‘conventional wisdom’ of early virulence evolution theory, speculates that pathogens should evolve over time towards avirulence or commensalism with the host, and that virulence is a reflection of a novel host-parasite association [8]. In contrast, the short-sighted evolution hypothesis postulates that pathogens evolve higher virulence in response to immediate within-host selection pressures, meanwhile sacrificing their long-term evolutionary advantage by harming the host [9]. The trade-off hypothesis predicts that pathogens will optimize their overall reproductive fitness by trading off between virulence and transmission, selecting for intermediate virulence [10]. Unlike the other models that look to within-host determinants, the coincidental selection hypothesis argues that virulence may be inconsequential to success in the host of interest, evolving in the environment or another host and merely maintained due to minimal impact on fitness [6]. While these models of virulence evolution have been studied extensively in a number of biological systems [10-12], there have been few empirical tests as to how virulence evolves in opportunistic pathogens during chronic infection [13].

Here, we tested how virulence evolves in an opportunistic pathogen during chronic infection, using the human opportunist, *P. aeruginosa*, in murine chronic wounds. *P. aeruginosa* is an ESKAPE pathogen notorious for multi-drug resistance [14] and a model organism for the study of chronic infections. It causes long-term infection in the lungs of cystic fibrosis (CF) patients and in chronic diabetic wounds [15]. *P. aeruginosa* is one of the most common bacterial pathogens isolated from chronic wounds, often forming antimicrobial-tolerant biofilms that are difficult to eradicate [16]. Chronic wounds present a massive burden on patients and healthcare systems worldwide, characterized by persistent infection, excessive inflammation, and a significantly delayed healing process [17-24]. While the adaptation of *P. aeruginosa* to CF lungs has been well-studied [25-28], long-term adaptation in chronic wounds is not as well-documented, presenting opportunities to study the nature of virulence evolution and pathogenesis in a clinically relevant environment.

Using a two-part serial passage selection and sepsis experiment, we determined how *P. aeruginosa* virulence evolves in murine chronic wounds, and whether the evolution of virulence was reproducible. We also ascertained morphological diversity and phenotypic changes after 42 days and ten rounds of evolution in wounds, and used whole genome sequencing to identify genetic signatures of *P. aeruginosa* adaptation to a chronic wound environment.

## MATERIALS & METHODS

### Bacterial Strains and Culture Conditions

We infected mice with the *Pseudomonas aeruginosa* strain PA14. For overnight cultures, we grew cells in 24-well microtiter plates in lysogeny broth (LB) and incubated at 37°C with shaking at 200 rpm.

### Serial passage experiment

The murine chronic wound model used in this study is based on one that has been previously described [29-31]. We anesthetized adult female Swiss Webster mice (Charles River Laboratories, Inc.), weighing between 20-25 g, by intraperitoneal injection of 100 mg/kg sodium pentobarbital (Nembutal; Diamondback Drugs), before their backs were shaved, and the hair was cleanly removed with a depilatory agent. As a preemptive analgesic, 0.05 mL of lidocaine (500 µL of bupivacaine [0.25% w/v] with 500 µL of lidocaine [2% w/v]) was injected subcutaneously in the area to be wounded. We then administered a dorsal 1.5 x 1.5 cm excisional skin wound to the level of the panniculus muscle, and covered it with transparent, semipermeable polyurethane dressings (OPSITE dressings) and injected approximately 10^3^ bacterial cells suspended in LB into the wound bed to establish infection. This adhesive dressing prevents contractile healing and ensures that these wounds heal by deposition of granulation tissue. At the end of the 72 h experimental infection period, we euthanized the animals and harvested their wound beds and spleens for colony forming unit (CFU) quantification on *Pseudomonas* Isolation Agar (PIA). We collected and saved a lawn of the 1:1000 dilution of each wound bed population in BHI + 25% v/v glycerol, then re-grew the cryo-stored population of the previous mouse in a new LB culture and inoculated, as before, into a new animal (Fig. S1A). We used three parallel groups of 10 mice (n=30 in total) to establish three independent evolution lines (A, B, and C), with the initial mouse of each group being inoculated with a stock population of PA14.

### Sepsis model

In a chronic wound model, sepsis is an important indicator of virulence, as septicemia is one of the most life-threatening outcomes of a chronic wound in human patients. From each of the 10^th^ and final evolved populations from the serial passage experiment, along with the ancestral PA14, we grew the cryo-stored wound populations in LB. We used each of these four liquid cultures to inoculate a distinct set of five mice with ∼10^5^ bacterial cells (n=20 in total; Fig. S1B). We monitored these mice for 80 hours for development of sepsis. If a mouse was moribund during this period, we euthanized it and harvested the spleen for CFU counts. At the end of 80 h, we euthanized all remaining mice and harvested spleens for CFU counts. Due to the spleen’s role in the host immune response and blood filtration during infection, it is often one of the first organs to become infected post-septicemia. As such, bacterial load in the spleen is a better indicator of systemic infection and more relevant when discussing virulence and host health, while wound bed bacterial load is primarily an indicator of infection severity at the site of infection [32]. Therefore, we chose to only measure spleen colony forming units (CFUs) for the sepsis experiment.

### Colony morphology

To assess the diversity in colony morphology of evolved populations, we plated serial dilutions of the previously cryo-stored populations of evolutionary rounds 5 and 10 on Congo red agar (CRA) plates [33]. We chose CRA to highlight any rare colony morphology types that may otherwise be overlooked on LB agar. We randomly picked 100 colonies from each of these populations to start overnight cultures, and from these, made cryo-stocks of each isolate and plated 1μL on CRA to compare individual colony morphologies. All CRA plates were incubated at 37°C overnight, then for 3-4 days at room temperature to allow for full development of colony morphologies.

### Whole genome sequencing and variant calling

We chose one representative isolate from each morphotype and line of evolution at round 10 in addition to the ancestral PA14 for sequencing analysis. We streaked the cryo-stocks of these isolates on LB agar and picked single colonies, from which we grew overnight cultures in LB broth. We isolated genomic DNA from the liquid cultures using the DNeasy Blood and Tissue Kit (Qiagen) according to the manufacturer’s instructions. We prepared sequencing libraries using the Nextera XT DNA Library Preparation Kit and sequenced with the Illumina Miseq platform, with a minimum average calculated level of coverage of 30× for each selected isolate. We first trimmed reads and removed adapter sequences, then mapped all samples against *P. aeruginosa* PA14 (RefSeq accession number GCF_000404265.1), and called single nucleotide polymorphisms (SNPs), insertions, and deletions using the reference-based alignment and variant calling tool *breseq* with default parameters. We manually parsed this list to eliminate any mutations erroneously called due to errors in sequencing alignment. Lastly, we determined the variants called between the reference PA14 and our ancestral PA14, then manually checked these against the list of indels and SNPs of all evolved isolates to create the final table of evolved mutations. We confirmed all mutations occurring in coding regions of defined proteins with Sanger sequencing.

### Pyocyanin assay

The pyocyanin assay is based on one that has been previously described [34]. We grew all isolates overnight in LB, then standardized OD_600_ of all cultures to 1.0 using phosphate-buffered saline (PBS). We spun cultures down briefly in a microcentrifuge before filtering through 0.2μm pore size syringe filters. We extracted 1mL of filter sterilized supernatant with 600μL chloroform, vortexed for 2 minutes, then centrifuged at 10,000 rpm for 5 minutes. We discarded the clear layer and re-extracted the blue layer with 400μL of 0.2M HCl, vortexed again for 2 minutes, and centrifuged at 10,000 rpm for 5 minutes. We then transferred the pink layer into a clear 96-well plate (Corning) and read the optical density at 520 nm.

### Pyoverdine and pyochelin production

Succinate media and siderophore production assay were modified from multiple sources [35-40]. Succinate media used for these assays was composed of 6g K_2_HPO_4_, 3g KH_2_PO_4_, 1g (NH_4_)_2_PO_4_, 0.2g MgSO_4_, and 4g succinic acid to a final volume of 1L H_2_O, pH adjusted to 7. We first grew all isolates overnight in LB, spun down 500μL of overnight LB culture, rinsed 2× with equal volume succinate media, and used this starter culture to inoculate 5mL of succinate media. We grew succinate cultures for 36 h at 30°C, as this medium and culture condition has been shown to maximize siderophore production [39], so as to highlight differences in production capability between isolates. We filtered cultures using 0.2μm pore size syringe filters, and transferred 100μL of supernatant to a black 96-well clear bottom microtiter plate (Corning). We measured pyoverdine fluorescence with an excitation wavelength of 400nm, emission wavelength of 460nm, and gain of 61. We measured pyochelin fluorescence with an excitation wavelength of 350nm, emission wavelength of 430nm, and gain of 82. We standardized all fluorescence values by the OD_600_ of each culture.

### Protease activity

We prepared skim milk agar plates composed of 5% w/v dry milk with 1.25% w/v agar. We poured 15mL of skim milk agar in 100 × 15mm Petri dishes. We grew liquid cultures overnight from a single colony in LB, then standardized OD_600_ of all cultures to 1.0 using PBS. We spun cultures down briefly in a microcentrifuge before filtering through 0.2μm pore size syringe filters. We spotted 10μL of filtered supernatant on skim milk agar plates, using 10μL of LB as a negative control and 1μL of proteinase K as a positive control. We incubated plates at 37°C overnight and measured the zone of protease activity qualitatively.

### Swarming motility

The components for swarm agar and experimental protocol were adapted from multiple sources [41-43]. Swarm agar was composed of 1× M8 salt solution (64g Na_2_HPO_4_ ·7H_2_O or 30g Na_2_HPO_4_, 15g KH_2_PO_4_, and 2.5g NaCl to a final volume of 1L H_2_O), 0.6% w/v agar, 0.5% w/v casamino acid, 0.2% w/v glucose, and 1mM MgSO_4_. We poured 25mL of swarm agar in 100 × 15mm Petri dishes under laminar flow, allowing for plates to dry for 30 minutes with plate lids off. We grew liquid cultures overnight from a single colony and inoculated plates with 2.5μL of overnight culture, incubating in short stacks of ≤4 plates, right side up for approximately 20 h.

### Swimming motility

Swim agar was composed of LB with 0.3% w/v agar. We poured 25mL of swim agar in 100 × 15mm Petri dishes, allowing a few hours to dry at room temperature with plate lids closed. We grew isolates overnight from a single colony, dipped a toothpick into the overnight culture, and inoculated by sticking the toothpick in the center of each plate, halfway through the agar. We wrapped short stacks of ≤4 plates in cellophane and incubated overnight for 20 h at 37°C alongside two large containers of water to retain humidity in the incubation chamber.

### Statistical analysis

We used a Kruskal-Wallis one -way test of variance to test for the difference of means, followed by a *post hoc* Dunn’s test with either a Holm-Bonferroni family-wide error rate (FWER) or Benjamini & Hochberg false discovery rate (FDR) correction. We used a Pearson’s correlation test to test the linear correlation between variables. Statistical significance was determined using a p-value < 0.05. We plotted graphs and performed statistical analysis in R version 3.6.1 using the packages tidyverse [44], ggplot2 [45], ggpubr [46], and PMCMR [47].

## RESULTS

### Wound bed and spleen bacterial population densities are positively correlated

We assessed the changes in bacterial load during the course of selection and found that wound bed CFUs throughout the serial passage experiment were generally within two orders of magnitude, aside from one mouse in evolutionary line A at round 8, whose bacterial load was notably lower (Fig. 1A). The bacterial load in spleens was highly variable across all three replicate lines of evolution, with many values being below our limit of detection, as the lowest serial dilution we plated was 10^−2^ (Fig. 1B). There was a positive correlation between bacterial load in wound bed and spleen during the serial passage experiment (Pearson’s r(28) = .44; *p* = 0.015).

**Figure 1.**
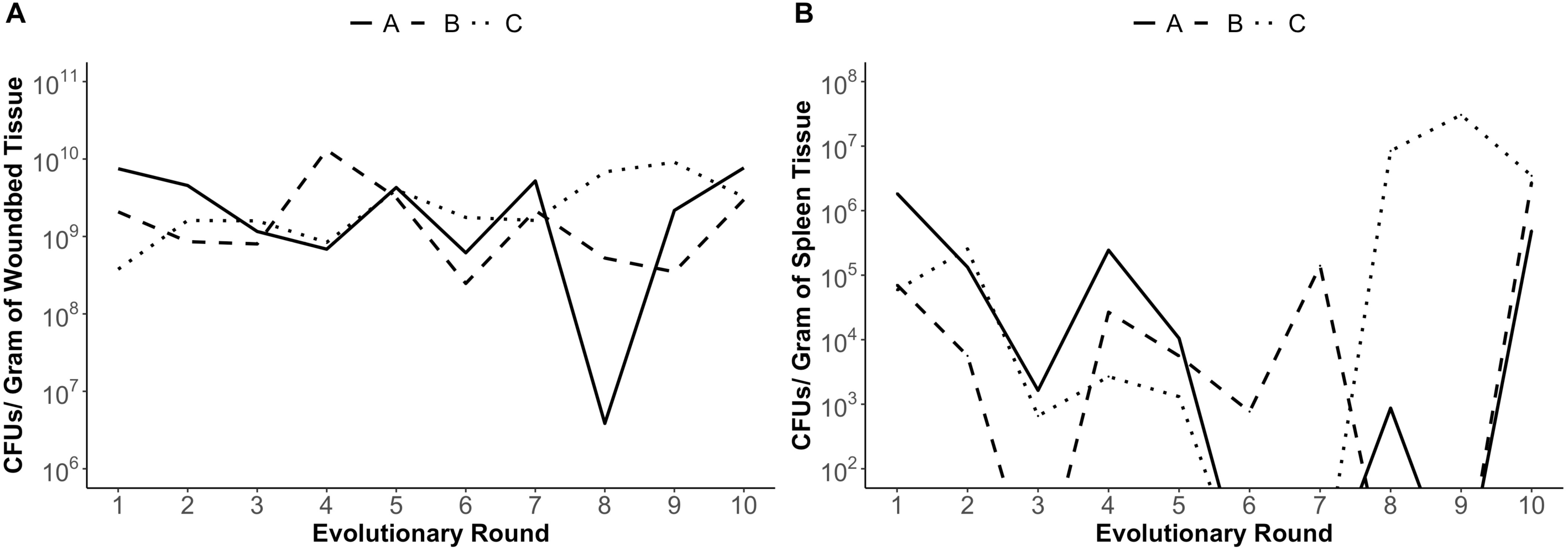
*P. aeruginosa* population densities in wound bed and spleen tissues during serial passage experiment are positively correlated. (A) Wound bed CFUs for mice at time of death for each evolutionary round were relatively stable, aside from the 8^th^ mouse in line A. (B) Spleen CFUs for mice at time of death for each evolutionary round were highly variable throughout the experiment, with many values falling below our limit of detection (10^2^ cells). Each CFU count represents one technical replicate. Wound bed and spleen CFUs during the serial passage experiment were positively correlated (Pearson’s r(28) = .44, *p* = 0.015).

### Morphological diversity is limited in chronic wound adapted populations

As diversity has been extensively reported in cystic fibrosis (CF) infections of *P. aeruginosa* [25-27, 48-50], we therefore characterized *P. aeruginosa* adaptation to chronic wounds and assessed population heterogeneity after 42 days of evolution. We began by characterizing the morphology of 100 random isolates from populations of rounds 5 and 10 of each evolutionary line. We assessed the diversity in colony morphology types (morphotypes) using Congo red agar plates. At round 5, each evolutionary line contained only 1-2 distinguishable morphotypes. At round 10, line A had two distinguishable morphotypes, while lines B and C each had three (Fig. 2A; Table S1). Isolates are named for their evolutionary line and the order in which they were characterized. We chose one representative isolate from each morphotype and line of evolution (A88, A92, B16, B31, B42, C31, C38 and C62) for further analysis.

**Figure 2.**
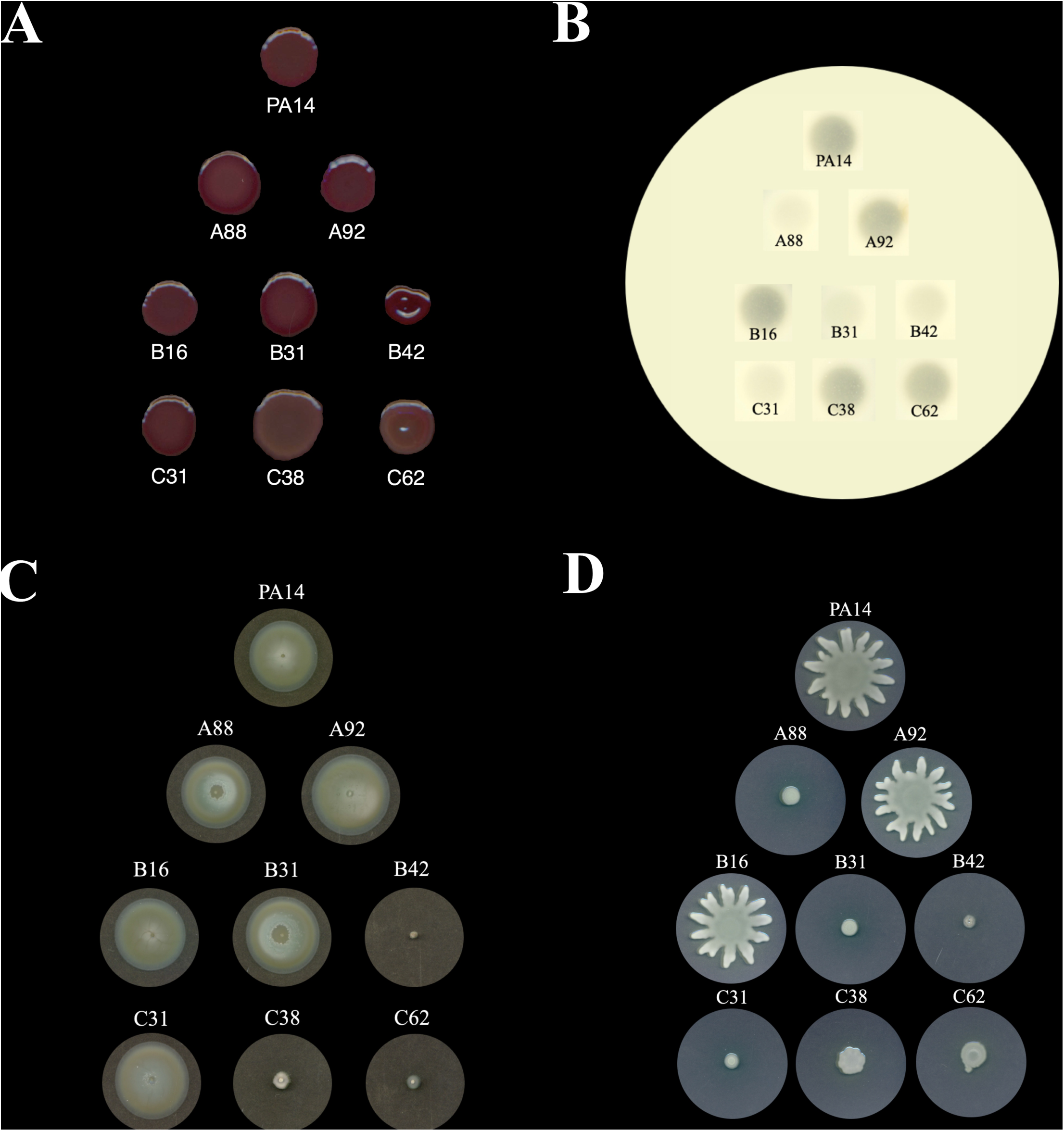
Changes in morphology, protease production, and swimming and swarming motilities. (A) There were five distinct types of colony morphology on CRA at the final round of selection across all three lines of evolution, with line A having two distinct morphology types, and lines B and C each having three distinct colony morphology types (with some colony morphology types being present in multiple lines). (B) Isolates A92, B16, C38, and C62 displayed protease activity comparable to that of the ancestral PA14, while isolates A88, B31, B42, and C31 showed decreased protease activity. (C) Isolates B42, C38, and C62 lost the ability to swim. (D) Isolates A88, B31, B42, C31, C38, and C62 lost the ability to swarm.

We tested these representative isolates for total protease activity, pyoverdine, pyochelin, and pyocyanin production, and swimming and swarming motility, to assess any phenotypic variation potentially relevant to *P. aeruginosa* pathogenesis between the evolved isolates and the ancestor [15]. Isolates A92, B16, C38, and C62 produced similar levels of protease production to PA14; while A88, B31, B42, and C31 demonstrated decreased relative protease activity (Fig. 2B). A88, A92, B16, B31, and C31 displayed swimming motility; however, only A92 and B16 showed fully functioning swarming motility (Fig. 2B-C). There were differences in pyoverdine (χ^2^(8) = 20.186, *p* = 0.0097), pyochelin (χ^2^(8) = 19.17, *p* = 0.014), and pyocyanin (χ^2^(8) = 47.843, *p* = 1.059e-7) production between isolates (Fig. 3).

**Figure 3.**
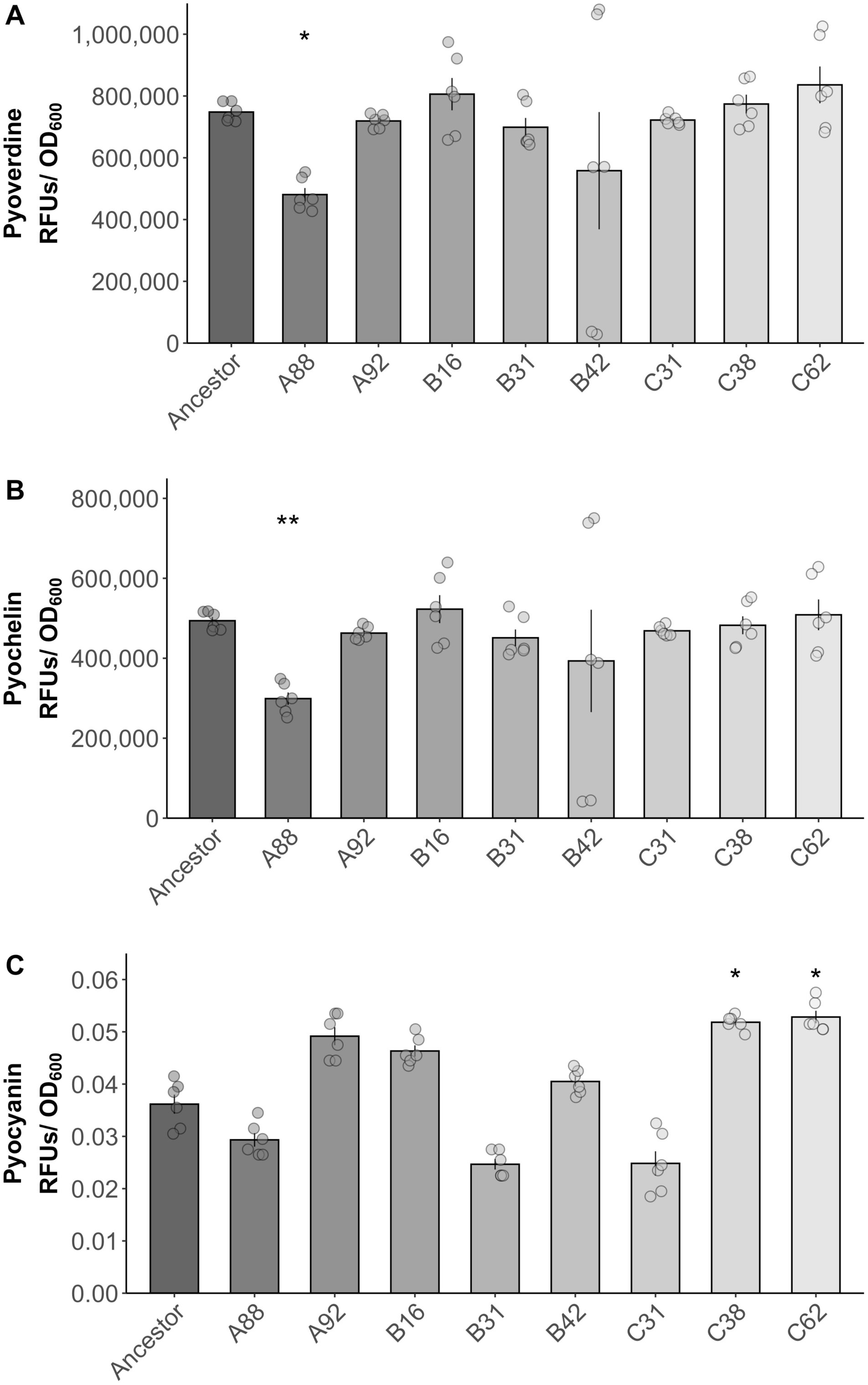
Changes in production of pyoverdine, pyochelin, and pyocyanin. (A) Pyoverdine production in the final evolved representative isolates and ancestral PA14. A88 was the only evolved isolate with pyoverdine production significantly different than the ancestor strain (Kruskal-Wallis, Dunn’s post hoc test, Benjamini & Hochberg correction, *p* = 0.018). Error bars indicate SEM. (B) Pyochelin production in the final evolved representative isolates and ancestral PA14. A88 was the only evolved isolate with pyochelin production significantly different than the ancestor strain (Kruskal-Wallis, Dunn’s post hoc test, Benjamini & Hochberg correction, *p* = 0.0082). (C) Pyocyanin production in the final evolved representative isolates and ancestral PA14. C38 and C62 both displayed pyocyanin production significantly different than the ancestor strain (Kruskal-Wallis, Dunn’s post hoc test, Benjamini & Hochberg correction, *p* = 0.01975 and *p* = 0.01915, respectively).

### Genes encoding virulence determinants are mutated during evolution in a chronic wound

Loss of virulence factors and social traits through genetic mutations is commonly observed in *P. aeruginosa* isolates collected from chronic CF infections [51-53]; however, the genetic adaptations of *P. aeruginosa* to chronic wounds are less well described. We conducted whole genome sequencing on each of the representative morphotypes to identify possible genetic signatures of adaptation to chronic wounds. Only a small number of mutations were identified, some occurring across more than one line of evolution. Across all three lines, we found in total seven unique mutations, six of them resulting in a change in amino acid sequence within a coding region (Table 1). We identified mutations in *lasR, pvcA, fleQ, rpoN, pilR*, all genes previously implicated in *P. aeruginosa* virulence [54-65]. The same *lasR* and *pvcA* mutations were found in each of the three evolutionary lines. There was additionally one frameshift mutation located within a coding region for a hypothetical protein with no known homologs.

**Table 1.** A list of all mutations in the final evolved representative morphology type isolates as mapped to the PA14 reference genome. Many genes coding for virulence factors or regulators of virulence are mutated over the course of adaptation to chronic wounds.

| Gene locus | Gene annotation | Genetic mutation | Amino acid effect | Isolate(s) |
| --- | --- | --- | --- | --- |
| PA14_33290/<br>PA14_33300 | Intergenic region | Δ180 bp | N/A | C62 |
| PA14_35430 | *pvcA* | GCG → ACG | A249T | A88, B31, B42, C31 |
| PA14_45960 | *lasR* | Δ9 bp at pos. 130-138 | ΔS44-D46 | A88, B31, B42, C31 |
| PA14_50220 | *fleQ* | GTC → GGC | V270G | C38, C62 |
| PA14_57940 | *rpoN* | GAC → AAC | D459N | B42 |
| PA14_60260 | *pilR* | ACC → CCC | T275P | C62 |
| PA14_70360 | Hypothetical protein | Δ36 bp at pos. 98-133 | Frameshift mutation | C38, C62 |

### Virulence can evolve divergently in chronic wounds

To assess how virulence evolved in our wound model, we compared the virulence of each of the three evolved populations against each other and the ancestral PA14 using a sepsis experiment (Fig. S1B). We observed that at the end of the sepsis experiment, three of the five mice infected by evolution line C survived, two from the ancestral PA14, one from line A, and none from line B (Fig. 4A). A Kruskal-Wallis test showed that there were significant differences in the mean spleen CFUs at the time of death between mice infected by the various populations (χ^2^(3) = 10.623, *p* = 0.014; Fig. 4B). A *post hoc* analysis showed that this statistically significant difference was between mice infected by lines B and C (*p* = 0.023, Dunn’s test, Holm-Bonferroni correction). Mice infected by the ancestral PA14 and line B showed differences in spleen CFUs, just above the α = 0.05 significance threshold (*p* = 0.058). Overall, we found that over the course of 42 days, line B evolved to be more virulent, line C evolved to be slightly less virulent, and line A remained approximately as virulent as the ancestor.

**Figure 4.**
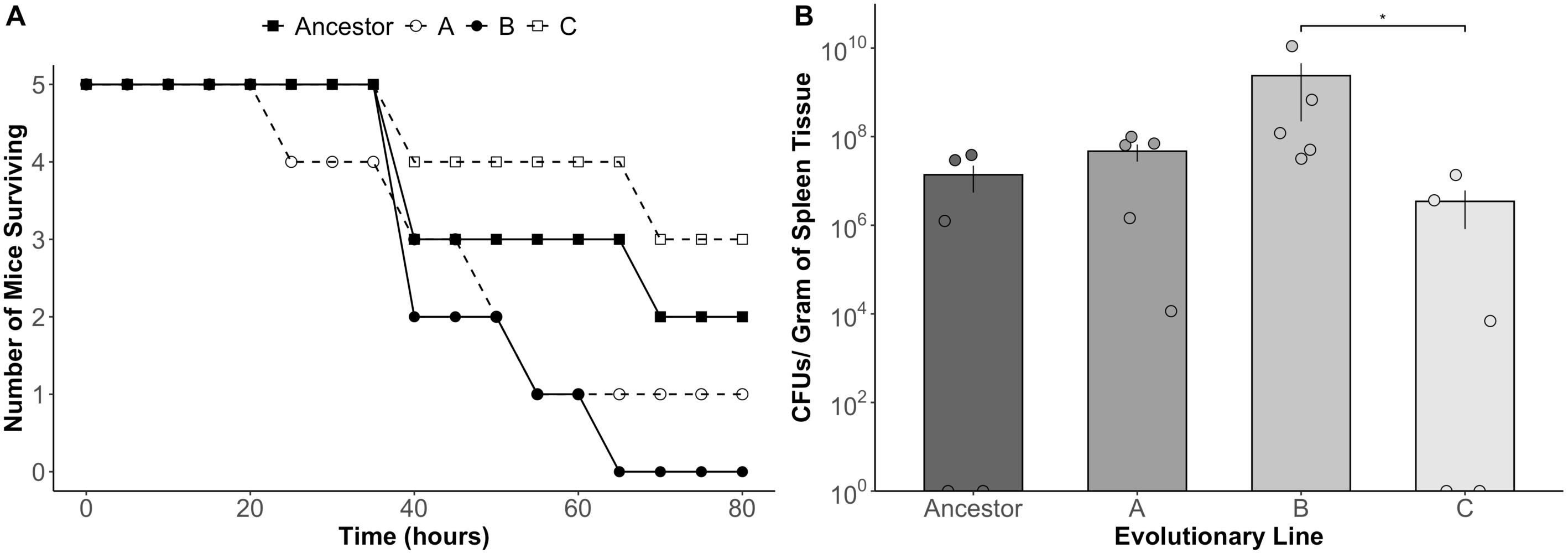
Virulence can evolve in diverging directions in a chronic wound. (A) Mice infected by the final evolved population of line B in the sepsis experiment had the highest mortality rate (100%), with no surviving mice at the end of 80 hours, while mice infected by line C had the lowest mortality (40%), with three of five mice surviving. (B) Mice infected by the final evolved population of line B in the sepsis experiment had significantly higher mean spleen CFUs at time of death as compared to mice infected by line C, indicating more severe septicemia (Kruskal-Wallis, Dunn’s *post hoc* test, Holm-Bonferroni correction, *p* = 0.023). Error bars indicate SEM.

## DISCUSSION

Opportunistic pathogens and their resulting chronic infections pose a significant healthcare burden, affecting 1-2% of the population in developed nations and amounting to billions of dollars annually in treatment costs [17-24]. Yet, the nature of virulence evolution in these organisms during chronic infection remains poorly understood. To assess how virulence evolves in an opportunistic pathogen during a chronic infection, we passaged *P. aeruginosa* PA14 in a murine chronic wound model for ten rounds of selection spanning 42 days, isolated a number of strains after the ten rounds for phenotypic and genotypic analysis, and compared the virulence of the whole evolved populations with that of the ancestor strain using a mouse sepsis experiment. We found that (i) there was a lower degree of morphological and phenotypic diversity in our evolved populations than has been previously described for *P. aeruginosa* in CF infections [27, 48, 49]; (ii) our populations acquired mutations in major regulators and genes previously shown to be involved in pathogenesis; and (iii) virulence evolved differently in each of the three independent evolutionary lines.

To interpret these results and make meaningful predictions for human chronic infections, we must consider the following caveats: (i) time scale of our experiment, (ii) growth in the environment, and (iii) the number of isolates tested for phenotypic and genotypic analysis. The time scale of our experiment, 42 days, is significantly shorter than that of a human chronic wound infection. Additionally, we conducted generations of growth in LB medium in between rounds of evolution in mice, which may have introduced another variable of selection. We also acknowledge that while the sepsis experiment is a measure of population-level virulence, the virulence factor phenotypes, i.e. proteases, siderophores, pyocyanin, and motility, were conducted on a small sub-sample of isolates and may not be reflective of the population-level phenotype. Given all of these considerations, caution must be exercised when extrapolating experimental results from a laboratory mouse model to a vastly more complex human infection, as virulence factors can be host specific [66].

We first assessed morphotypic diversity within *P. aeruginosa* populations after ten rounds of selection. Previous studies on *P. aeruginosa* in the CF lung have shown a high degree of heterogeneity, with populations displaying up to 15 different morphotypes in a single patient’s sputum sample [67, 68]. In our study, we only identified five distinct morphotypes which were smooth, non-mucoid, of similar size and with only small variations in pigment production. The low degree of morphotypic diversity we observed may be due in part to the time scale of our experiment, as six weeks is not comparable to the years of evolution in a CF lung. In addition, a chronic wound may lack the spatial structure seen in CF lungs [50], providing fewer niches for diversification. Further, our experiment focused on a monospecies infection, and CF lungs (and human chronic wounds) are comprised of polymicrobial infections, which may encourage further diversification through microbial interactions [69].

To understand how *P. aeruginosa* adapts to a chronic wound environment, we whole genome sequenced one isolate of each representative morphotype after ten rounds of selection. We identified mutations in *lasR, pvcA, rpoN, fleQ*, and *pilR*, all genes previously shown to be important for *P. aeruginosa* virulence [54-65]. The occurrence of the same *pvcA* and *lasR* mutations in all three lines presents evidence of parallel evolution, suggesting that these mutations may confer fitness advantages in wounds. LasR plays an important role in the quorum sensing (QS) hierarchy of *P. aeruginosa*, acting as a transcriptional activator for a plethora of genes implicated in virulence [57]. Although the in-frame deletion we observed in LasR, ΔS^44^-D^46^, is not at an active site, it is in extremely close proximity to a number of residues forming a ligand-binding pocket— G^38^, L^39^, L^40^, Y^47^, E^48^, and A^50^ [70]. Furthermore, it is conceivable that a three-residue deletion could significantly impact protein folding and lead to a loss-of or decreased function. This is in agreement with the phenotypes we observed, as our *lasR* mutants showed decreased protease production and inhibited swarming ability, and it has been previously shown that *lasR* mutants show diminished swarming behavior [41]. PvcA is involved in the biosynthesis of paerucumarin, a molecule which has been suggested to enhance the expression of iron-regulated genes by chelating extracellular iron [64, 65]. The *pvcA* mutation we observed in all three lines (A^249^T) of a non-polar to a polar side-chain amino acid, while not in an active site, could reasonably result in a detrimental impact to three-dimensional protein folding, and potential loss-of-function [71].

The mutations in *rpoN, fleQ*, and *pilR* all lead to amino acid changes within highly conserved domains or residues directly involved in the activity of their respective protein products [72-74]. The gene product of *rpoN*, RNA polymerase factor σ^54^, regulates a wide variety of functions in *P. aeruginosa*, including the *rhl* QS system, flagellin, and pilin production, which play important roles in motility, surface attachment, and biofilm formation [54, 55, 60-63]. Likewise, our *rpoN* mutant, B42, showed inhibited motility. FleQ is a transcriptional regulator for both flagellin and exopolysaccharide (EPS) biosynthesis in *P. aeruginosa* [58]. The *fleQ* mutants, C31 and C62, displayed diminished swimming and swarming motilities. Lastly, PilR is a transcriptional regulator for type IV pili expression, a structure involved in twitching motility and DNA uptake [56]. While we did not phenotypically assess twitching motility, a mutation in a core functional residue suggests most likely loss-of-function. The locations of these mutations, along with the supporting phenotypic data, suggest that all of these mutations have led to loss-of-function. It has previously been shown that in chronic CF infections, *P. aeruginosa* selects against the production of virulence factors that are required for acute infection [27, 53]. Many CF isolates are *lasR, rpoN*, and *fleQ* mutants [25-28]; likewise, another long-term evolution experiment of *P. aeruginosa* found an accumulation of *lasR* and various *pil* mutants [75]. Our results, taken with previous studies, suggest that *P. aeruginosa* may employ similar genetic adaptations in multiple infection environments.

We found that evolution, with respect to levels of overall virulence, was not reproducible in our three independent selection lines, in contrast to a previous study that showed *P. aeruginosa* evolution was highly reproducible *in vitro* [76]. This highlights the likely importance of host-specific variables such as the immune response on *P. aeruginosa* evolution and virulence. None of the current classic virulence models (see introduction) in isolation adequately explains this diverging pattern, suggesting there may be previously overlooked variables influencing virulence evolution in opportunistic pathogens, or that components from multiple virulence models may need to be considered in tandem. Heterogeneity within *P. aeruginosa* populations may partially account for the differing virulence trajectories we observed, as within-host adaptation leading to multiple infecting genotypes can result in higher or lower virulence, depending on the context. Levin & Bull originally proposed the short-sighted evolution hypothesis to explain the role of multiple infection and within-host selection on virulence [9]. According to their model, as a strain mutates and diversifies within the host, competition for limited resources will favor fast-growing genotypes, leading to an overall higher virulence. An alternative idea was proposed by Buckling and West, where virulence is predicted to decrease in response to heterogeneity or low relatedness, if virulence is dependent on the production of common goods and cooperation by members of the population [77]. Such common goods are exploitable by non-cooperating ‘cheats’ that increase their fitness in the population by benefitting from goods produced by cooperators, while undermining the pathogenicity of the population as a whole [78-82]. We found some support for both of these ideas, as we observed higher virulence in one evolutionary line but lower virulence in another.

In *P. aeruginosa*, quorum sensing (QS) controls the production of a number of secreted common goods, and strains isolated from chronic infections frequently display mutations in the QS regulator *lasR*. We identified *lasR* mutations in four of our isolates, but it remains to be determined whether they arose via social cheating or because they are better adapted to a wound environment.

However, their presence in our evolved populations is a plausible explanation for the reduction in virulence in one selection line, as the frequency of *lasR* mutants in populations has previously been linked to a reduction in social traits, virulence and antibiotic resistance [76, 79, 80, 82]. Another study on the evolution of *P. aeruginosa* in *Caenorhabditis elegans* also identified that virulence evolution was mediated by the production of secreted molecules. and that attenuation in pathogenicity could be attributed to cheats exploiting the goods produced by cooperators [75]. Taken together, this highlights the importance of social interactions during chronic infection, and that heterogenous populations likely result in complex interactions that can impact overall community function [76, 83]. Future work in this area could focus on exploring genetic diversity in infections using deep sequencing, which may provide insights as to how allelic polymorphism and genetic heterogeneity impacts community function and the outcome of virulence [76].

Overall, our study adds to the breadth of knowledge on *P. aeruginosa* adaptations *in vivo*, showing that while *P. aeruginosa* employs similar adaptive strategies, i.e. loss of virulence factors, in both chronic wounds and CF lungs, heterogeneity in chronic wounds may be less extensive. Our findings also emphasize that more work needs to be performed to increase our understanding of the dynamics and drivers of virulence evolution in opportunistic pathogens during chronic infection. This is an important consideration given the increasing interest in developing anti-virulence management strategies.

## Data accessibility

All sequences have been uploaded to the NCBI SRA database (accession number PRJNA643594). Raw data, code, and Sanger sequencing results have been made available in the Dryad Digital Repository (https://doi.org/10.5061/dryad.000000021).

## Ethical statement

All animals were treated humanely and in accordance with protocol 07044 approved by the Institutional Animal Care and Use Committee at Texas Tech University Health Sciences Center in Lubbock, TX.

## Author contributions

SPD and KPR designed the study. JV and DF performed the experimental work and analyzed the data. All authors contributed to the writing of the manuscript.

## Funding and acknowledgements

This material is based upon work supported by the National Science Foundation Graduate Research Fellowship (Grant No. DGE-1650044) to JV; The Cystic Fibrosis Foundation (DIGGLE18I0) to SPD; Cystic Fibrosis Foundation (AZIMI18F0) to SA; CF@lanta (3206AXB to SA; National Institutes of Health (R21 AI137462-01A1) to KPR; Ted Nash Long Life Foundation to KPR; and Novo Nordisk Foundation (NNF17OC0025014) to UT. We wish to acknowledge the core facilities at the Parker H. Petit Institute for Bioengineering and Bioscience at the Georgia Institute of Technology for the use of their shared equipment, services, and expertise. We thank Sam Brown for comments on the manuscript.

## Supplemental Figure legends

**Figure S1.**
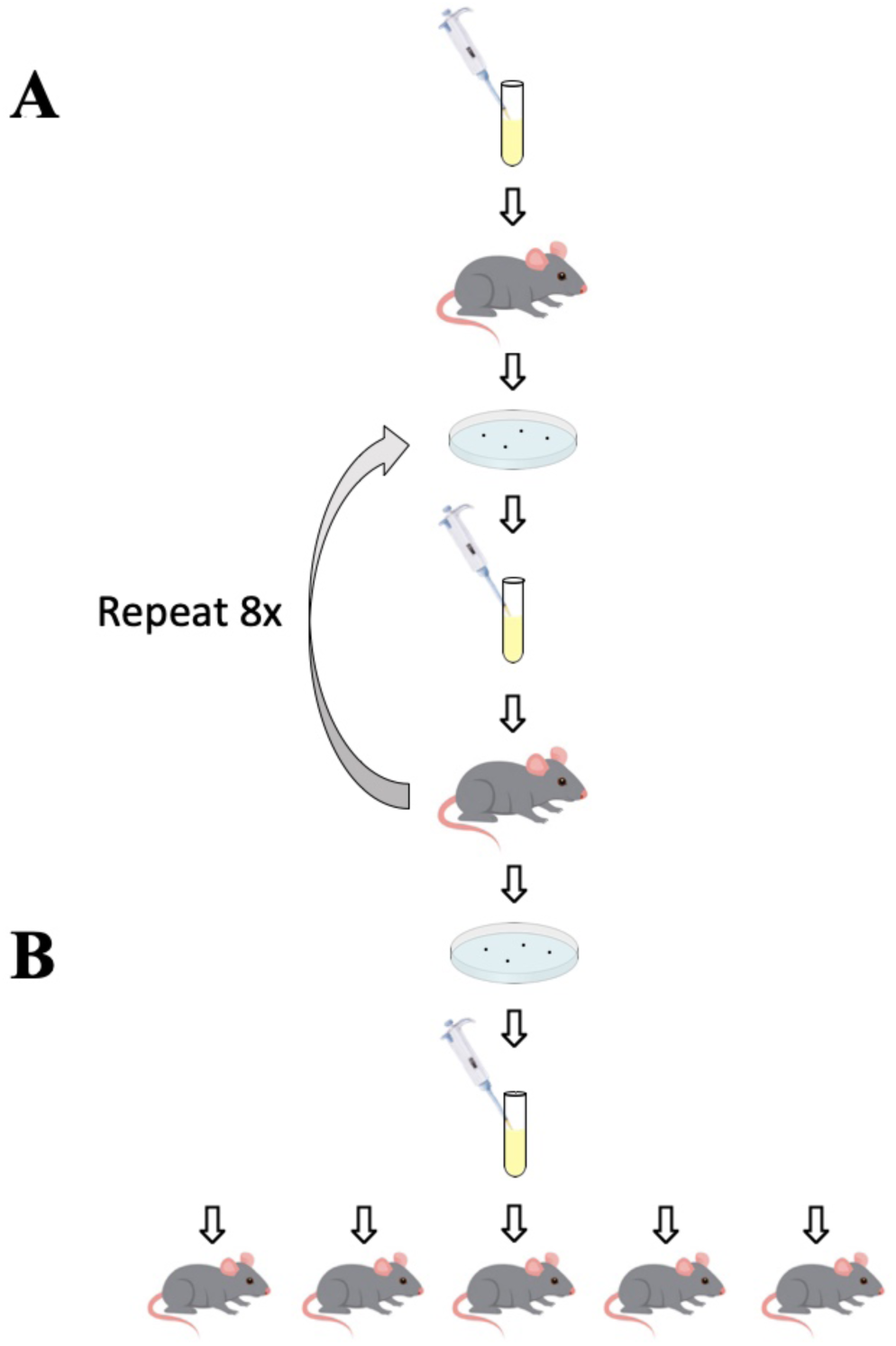
Serial passage and sepsis experimental protocols. (A) Serial passage experimental protocol. We established three independent evolution lines by infecting three mice with ∼10^3^ cells of the *P. aeruginosa* strain PA14 from a liquid lysogeny broth (LB) culture. Each infection duration was 72 h, after which we euthanized the mice and harvested their wound bed and spleen for colony forming unit (CFU) counts on *Pseudomonas* isolation agar (PIA). We used a 1:1000 serial dilution of the wound bed infection to start a new LB liquid culture and inoculate the next mouse in each line of evolution, again with ∼10^3^ cells. We carried this selection experiment through a total of ten passages in mice for each of the three parallel evolution lines (n=30 mice in total). One replicate evolutionary line is shown in the diagram. (B) Sepsis experimental protocol. We began the sepsis experiment by growing LB liquid cultures of the three final evolved populations from the serial passage experiment and of the ancestral PA14. We used each of these four liquid cultures to inoculate a distinct set of five mice with ∼10^5^ cells (n=20 mice in total). We monitored these mice for 80 h for sepsis. If a mouse was moribund, it was euthanized, time of death noted, and spleen harvested for CFU counts. At the end of 80 h, all remaining mice were euthanized, and their spleens harvested for CFU counts.

**Table S1.** Details of the biofilm colony morphology types of evolved isolates on CRA after ten rounds of selection spanning forty-two days across all three lines of evolution. One isolate of each distinct colony morphology type from each population was selected as a representative for further phenotypic and genomic analysis.

| Representative Isolate | Morphology type | Morphology Description |
| --- | --- | --- |
| PA14 | 1 | dull red, wrinkly edge, smooth texture |
| A88 | 2 | blood red, smooth edge, smooth texture |
| A92 | 1 | dull red, wrinkly edge, smooth texture |
| B16 | 1 | dull red, wrinkly edge, smooth texture |
| B31 | 2 | blood red, smooth edge, smooth texture |
| B42 | 3 | blood red, raised outer ring |
| C31 | 4 | blood red, smooth edge, smooth texture |
| C38 | 1 | dull red, wrinkly edge, smooth texture |
| C62 | 5 | dull red, smooth edge, smooth texture, faint outer ring |

